# Stabilising and destabilising kinesin complexes queue at plus tips to ensure microtubule catastrophe at cell ends

**DOI:** 10.1101/287649

**Authors:** John C. Meadows, Liam J. Messin, Anton Kamnev, Theresa C. Lancaster, Mohan K. Balasubramanian, Robert A. Cross, Jonathan B.A. Millar

**Affiliations:** Centre for Mechanochemical Cell Biology, Division of Biomedical Sciences, Warwick Medical School, University of Warwick, Gibbet Hill, Coventry CV4 7AL, UK

**Author notes:** These authors contributed equally to this work (John C. Meadows, Liam J. Messin).

## Abstract

In fission yeast, the length of interphase microtubule (iMT) arrays are adapted to cell length so as to maintain cell polarity and to help centre the nucleus and cell division ring. Here we show that length regulation of iMTs is dictated by spatially-regulated competition between MT-stabilising Tea2/Tip1/Mal3 (Kinesin-7) and MT-destabilising Klp5/Klp6/Mcp1 (Kinesin-8) complexes at iMT plus tips. During MT growth, the Tea2/Tip1/Mal3 complex remains bound to the plus tips of iMT bundles and restricts access to the plus tips by Klp5/Klp6/Mcp1, which accumulates behind it. At cell ends, Klp5/Klp6/Mcp1 invades the space occupied by the Tea2/Tip1/Tea1 kinesin complex triggering its displacement from iMT plus tips and MT catastrophe. These data show that *in vivo*, whilst the “antenna model” for iMT length- and age-dependent catastrophase accumulation has validity, length control is an emergent property reflecting spatially-regulated competition between multiple complexes at the MT plus tip.

## Introduction

Microtubule (MT) length control is important for multiple cellular processes including vesicle transport, mitotic spindle size, chromosome bi-orientation and cilia function (Kuan & Betterton, 2013; Niwa et al., 2012; Wang et al., 2016; Wordeman & Stumpff, 2009). In the fission yeast, *Schizosaccharomyces pombe*, arrays of interphase microtubules (iMT) grow along the long axis of the cell and undergo catastrophe at cell ends. Interaction of iMTs with the cell end cortex is required to maintain cell polarity and correctly position the nucleus and cell division ring (Huang et al., 2007; Martin, 2009; Tran et al., 2001). The maintenance of cell polarity and positioning of the division ring requires transport of the Kelch-repeat protein Tea1 and SH3-domain protein Tea4 on the plus tips of iMTs by association with Tea2 (Kinesin-7) and their deposition at cell ends following interaction of iMTs with the cell end cortex (Martin, 2009; Martin et al., 2005; Mata & Nurse, 1997; Sawin & Snaith, 2004; Snaith et al., 2005; Tatebe et al., 2005). Reconstitution and live-cell imaging experiments reveal that association of Tea2 (Kinesin-7) to the growing plus tips of MTs requires two other components, Mal3 (EB1 homologue) and Tip1 (Clip170 homologue) (Akhmanova & Steinmetz, 2008; Bieling et al., 2007; Busch & Brunner, 2004; Busch et al., 2004). The presence of the Tea2/Tip1/Mal3 complex, but not its cargo, at the MT plus tip also prevents premature MT catastrophe in the cytoplasm (Verde et al., 1995). Members of the Kinesin-8 family have attracted particular attention as regulators of MT length because they are both highly processive motors and undergo a conformational switch at the MT plus tip that promotes MT disassembly (Arellano-Santoyo et al., 2017; Su et al., 2012; Su et al., 2011; Tischer et al., 2009; Varga et al., 2006; Varga et al., 2009). These features have given rise to the “antenna model” for MT length control whereby more Kinesin-8 accumulates at the plus tips of longer MTs, thus increasing the likelihood of catastrophe and MT shrinkage (Leduc et al., 2012; Varga et al., 2006; Varga et al., 2009). Fission yeast contains two Kinesin-8 motors, Klp5 and Klp6, that function as an obligate heterodimer. Deletion of either gene causes numerous mitotic defects and overgrowth of interphase MTs, which result in defective nuclear positioning and loss of cell polarity, particularly in longer cells (Garcia et al., 2002; Grissom et al., 2009; Sanchez-Perez et al., 2005; West et al., 2002; West et al., 2001). Timely shrinkage of iMTs also requires Mcp1, a +TIP that is distantly related to the Ase1/PRC1/MAP65 family of anti-parallel MT binding proteins (Akhmanova & Steinmetz, 2010; Pardo & Nurse, 2005; Zheng et al., 2014). However, the precise role and relationship of these factors in controlling interphase MT length is unknown.

## Results and Discussion

We find that, like Klp5 and Klp6, the intensity of Mcp1 increases at plus tips of iMTs as they grow and dwell at cell ends and decreases as iMTs undergo shrinkage (Figure 1A & Figure 1 - figure supplement 1A). Importantly, binding of Mcp1 to iMT plus tips requires the motor activity of Klp5/Klp6 indicating that Mcp1 is a cargo of the Klp5/Klp6 complex (Figure 1 - figure supplement 1B-C; Zheng et al., 2014). Indeed, we find that Mcp1 binds weakly to Klp5 in co-immunoprecipitates from cell extracts (Figure 1 - figure supplement 1D). Importantly, the dwell time of iMTs at the cell end is extended in the absence of both Klp5 and Klp6 to the same extent as in the absence of Mcp1 and this effect is not additive, indicating that Mcp1 controls destabilisation of iMTs *via* its association with the Klp5/Klp6 complex (Figure 1B). Notably though, unlike deletion of either Klp5 or Klp6, loss of Mcp1 does not cause cell polarity defects in elongated *cdc25-22* cells (Figure 1 - figure supplement 2: West et al., 2001) and does not influence mitotic timing or accuracy of chromosome segregation (Figure 1 – figure supplement 3A-E). These functions may instead be due to association of Klp5/Klp6 to PP1, type-1-phosphatase (Dis2) (Meadows et al., 2011). Consistently, Mcp1 is not required for Klp5 and Klp6 to bind the mitotic spindle or kinetochores during mitosis and is not present in the nucleus during mitosis (Figure 1 - figure supplement 3F-G). We conclude, therefore, that Mcp1 is an interphase-specific regulator of Kinesin-8 mediated iMT length control in fission yeast.

**Figure 1.**
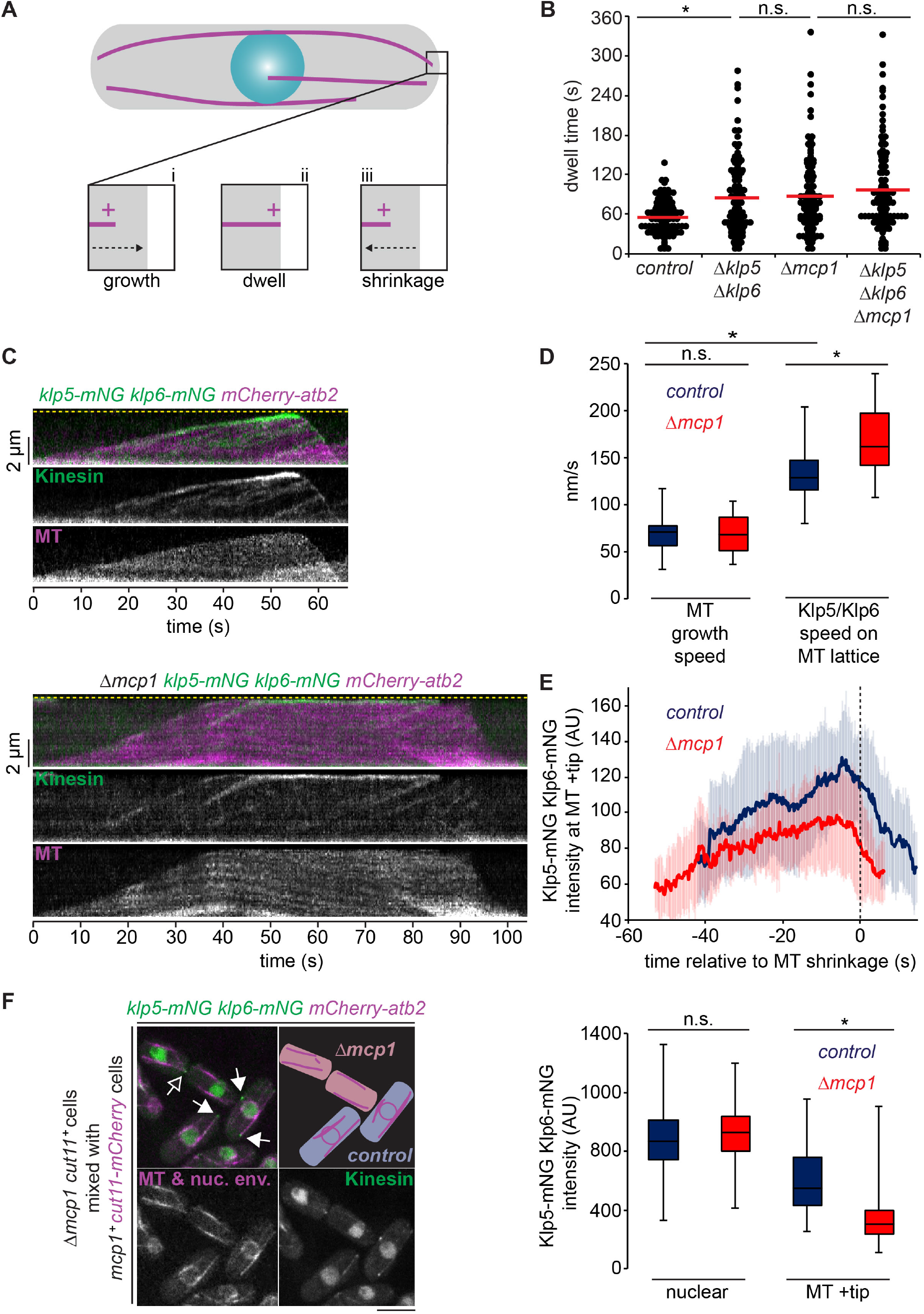
Mcp1 is required for control of interphase microtubule stability by Klp5/Klp6 but not for its motility. **(A)** Interphase microtubules (iMTs) (magenta) in fission yeast grow towards the cell end (i), dwell (ii) then shrink (iii). **(B)** Microtubules dwell at the cell end for the same extended period in cells deleted for *klp5* and *klp6*, *mcp1* or all three genes. Cells expressing fluorescently-tagged *α*2-tubulin (*atb2*) were imaged every 5 seconds and the dwell time of ∼100 individual iMTs recorded within the final 1.1 µm of the cell for each strain. Red bars signify the mean. Asterisk represents a p-value <0.001 and n.s. >0.05 (non-significant) between data sets calculated using the Kolmogorov-Smirnov test. **(C)** Klp5/Klp6 accumulates at the plus tips of both growing and dwelling microtubules. Kymographs showing fluorescently-tagged Klp5/Klp6 (Kinesin) co-imaged with fluorescently-tagged microtubules (MT) in the presence (top panels) and absence (bottom panels) of Mcp1. Dashed yellow line indicates the cell end. **(D)** Klp5/Klp6 speed exceeds iMT growth speed and is dependent on Mcp1. MT growth speed was calculated from kymographs from control (n=16) and *Δmcp1* cells (n=11), Klp5/Klp6 walk speed was calculated from multiple individual runs on the MT lattice in control (n=44) and *Δmcp1* cells (n=32). Asterisks represents p-values <0.001 and n.s. >0.05 (non-significant) between data sets calculated using the Kolmogorov-Smirnov test. **(E)** Average intensity of Klp5/Klp6 at the plus tips of iMTs from multiple kymographs of control (n=19) or *Δmcp1* cells (n=14). Error bars show standard deviation. **(F)** Mcp1 is required for full accumulation of Klp5/Klp6 at the iMT plus tip. Left panel shows the mixing experiment used to compare fluorescently-tagged Klp5/Klp6 levels between cells either expressing (blue, closed arrowheads) or deleted (red, open arrowhead) for Mcp1 in the same field of view. Controls cells can be distinguished from *Δmcp1* cells as they express a fluorescently-tagged nuclear envelope protein Cut11. Bar, 5µm. The box plot shows data quantitated from these experiments. Fluorescence values were recorded in the first frame for nuclear levels in control (n=44) and *Δmcp1* cells (n=45) and in the frame prior to MT shrinkage for levels at the MT plus tip in control (n=64) and *Δmcp1* cells (n=35). Measurements were normalised against levels of Klp5/Klp6 in the cytoplasm in all cases. Asterisk represents a p-value <0.001 and n.s. >0.05 (non-significant) between data sets calculated using the Kolmogorov-Smirnov test.

To further understand the role of Mcp1 in Kinesin-8 function, we examined Klp5/Klp6 motility and accumulation at iMT plus tips in control and *Δmcp1* cells by dual camera live-cell imaging (Figure 1C). We find that Klp5/Klp6 walks along the lattice of iMTs at 134 ± 28 nm/s, compared to average iMT growth speed of 69 ± 20 nm/s and accumulates at iMT plus ends, particularly as iMTs dwell at the cell end, and then dissociates following MT catastrophe (Figure 1C-E). In fact, Klp5/Klp6 walks faster (168 ± 37 nm/s) along the lattice of iMTs in the absence of Mcp1, although the growth speed of iMTs is unaffected. These data indicate that Mcp1 is required for the iMT destabilising function of Klp5/Klp6, perhaps by promoting its association to curved tubulin, but not for its processive motility. Instead it is likely that, like KIP3 and KIF18A, the highly processive motility of Klp5/Klp6 depends on MT binding site(s) in the C-terminal tail of those proteins (Mayr et al., 2011; Stumpff et al., 2011; Su et al., 2011; Weaver et al., 2011). To quantify the intensity of Klp5/Klp6 at plus tips, mixed populations of cells lacking Mcp1 and control cells were imaged in the same field of view. Although the absence of Mcp1 does not influence Klp5 or Klp6 stability, as judged by western blot of whole cell extracts (Figure 1 - figure supplement 1E), nor its intensity in the nucleus, Klp5/Klp6 accumulated to approximately half the level at iMT plus tips in the absence of Mcp1, even though iMTs dwell for longer at cell ends (Figure 1F). This may reflect a role for Mcp1 in enhancing the affinity of Klp5/Klp6 for the iMT lattice, resulting in additional runs to plus tips, or a role for Mcp1 in either retaining Klp5/Klp6 at plus tips or controlling the oligomeric status of the Klp5/Klp6 complex. Further work is needed to distinguish between these possibilities.

We next examined the functional relationship between the Klp5/Klp6/Mcp1 and Tea2/Tip1/Mal3 kinesin complexes. In contrast to Klp5/Klp6/Mcp1, which accumulates in both a MT length- and dwell time-dependent manner, binding of Tea2 kinesin to plus tips is independent of MT length (Figure 2A) but, like Klp5/Klp6/Mcp1, Tea2 dissociates from plus tips following MT catastrophe. To our surprise, we find that Tea2 remains bound, at the same average intensity, to stalled MT plus tips in the absence of Klp6 or Mcp1 (Figure 2B-C & Figure 2 – figure supplement 1A-B), indicating that the Klp5/Klp6/Mcp1 complex is required for timely dissociation of the Tea2 complex from plus tips. In the absence of Tea2, iMTs dwell at cell ends for less time than in control cells and undergo frequent MT catastrophe in the cytoplasm before iMTs reach the cell end, as previously observed (Figure 2D-F: Verde et al., 1995). Importantly, we find that deletion of *klp5* and *klp6* or *mcp1* in *Δtea2* cells largely restores MT catastrophe near the cell end and increases MT dwell time, although not quite to that observed in control cells (Figure 2E-F). A similar effect is seen in the absence of Tip1, but not in the absence of Tea1 (Figure 2 - figure supplement 2). This suggests that, in the absence of the Tea2/Tip1/Mal3 complex, but not its cargo, Klp5/Klp6/Mcp1 induces premature MT catastrophe in the cytoplasm. Consistently, Klp5/Klp6 accumulates at iMT plus tips in the absence of Tea2, albeit to a lower intensity than in control cells, as iMTs rarely last long enough to reach the cell end (Figure 2 - figure supplement 3A-B). In the absence of both Tea2 and Mcp1, Klp5/Klp6 accumulates at the plus tips of iMTs that reach the cell end, but at greatly reduced levels compared to control cells (Figure 2 - figure supplement 3C-D). In this situation, iMT catastrophe may purely be reliant on the force exerted by interaction of the plus tip with the cortex at the cell end (Tischer et al., 2009).

**Figure 2.**
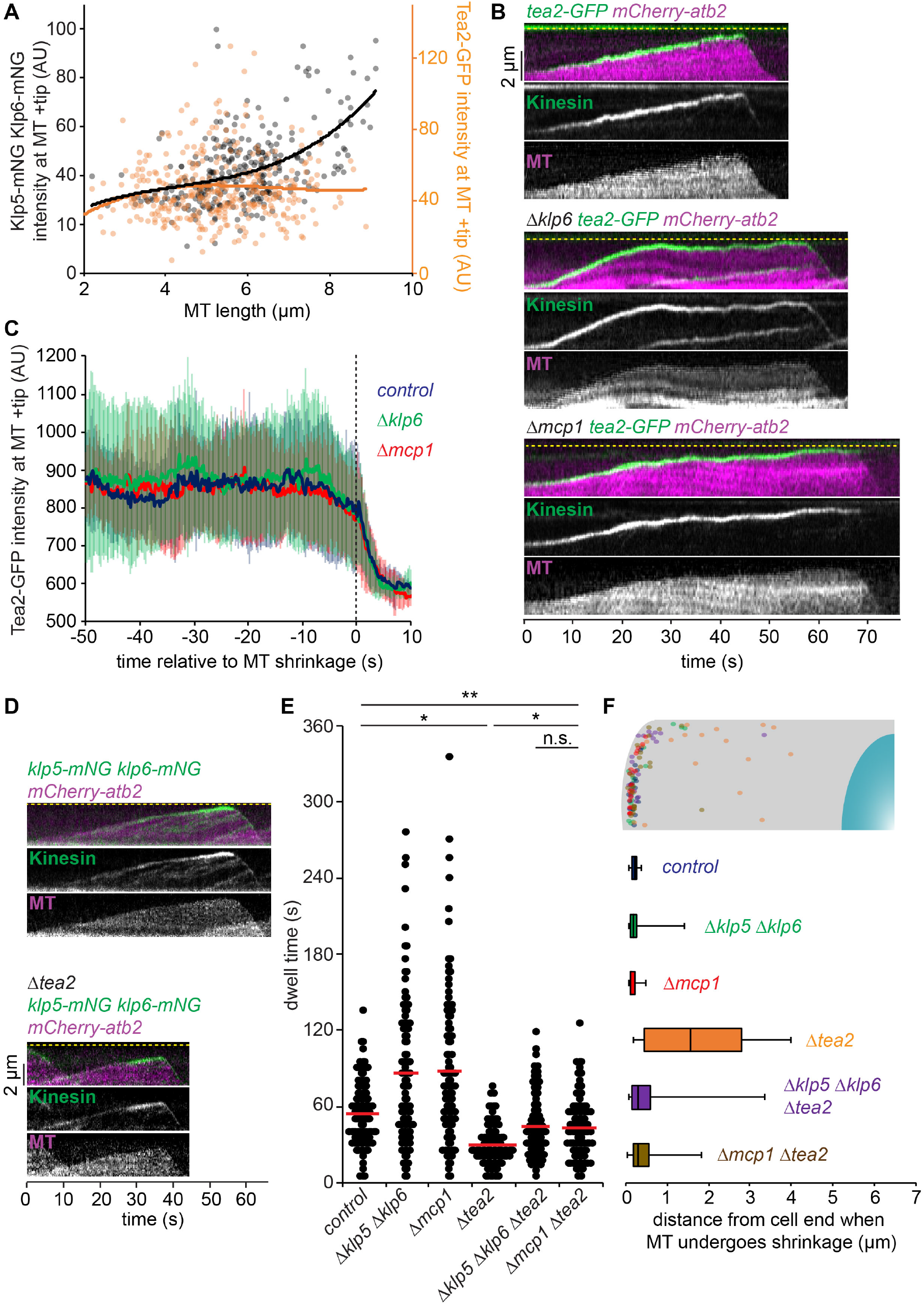
Tea2 and Klp5/Klp6/Mcp1 antagonistically control microtubule stability. **(A)** Klp5/Klp6 accumulates in a non-linear manner at the plus tips of growing MTs whereas Tea2 levels remain constant. Log phase cells expressing fluorescently-tagged MTs were imaged and the levels of either fluorescently-tagged Klp5/Klp6 or Tea2 at the plus tips of growing MTs determined. Measurements (*klp5-mNG klp6-mNG*, n=305; *tea2-GFP*, n=359) were plotted against microtubule (MT) length and third order polynomial curves fitted to the data. **(B)** Tea2 is retained at the plus tips of MTs that dwell for longer at the cell end in the absence of Klp6 or Mcp1. Kymographs showing fluorescently-tagged MTs co-imaged with fluorescently-tagged Tea2 kinesin in control cells (top panels) or in the absence of Klp6 (middle panels) or Mcp1 (bottom panels). Dashed yellow line indicates the cell end. **(C)** Intensity of Tea2 at plus tips quantitated from multiple kymographs of the type in (B) for control (n=16), *Δklp6* (n=13) or *Δmcp1* (n=14) cells (B). Error bars show standard deviation. **(D)** In the absence of Tea2, Klp5/Klp6 accumulates at plus tips before microtubules reach the cell end. Kymographs showing fluorescently-tagged MTs co-imaged with fluorescently-tagged Klp5/Klp6 in control (top panels) or *Δtea2* cells (bottom panels). Dashed yellow line indicates the cell end. **(E)** Antagonistic effect of Klp5/Klp6/Mcp1 and Tea2 on microtubule dwell time. Cells expressing fluorescently-tagged tubulin were imaged every 5 seconds and the dwell time of ∼100 individual iMTs for each condition recorded within the final 1.1 µm of the cell. Red bars signify the mean. Single asterisks represent p-values <0.001, double asterisks <0.05 and n.s. >0.05 (non-significant) between data sets calculated using the Kolmogorov-Smirnov test. **(F)** In the absence of Tea2 premature microtubule catastrophe in the cytoplasm is largely dependent on Klp5/Klp6/Mcp1. The strains in (E) were analysed to determine the position of 20 MT shrinkage events relative to the cell end. These events are represented both graphically and as a box plot.

We next considered how the antagonistic activities of the MT-stabilising Tea2/Tip1/Mal3 and MT-destabilising Klp5/Klp6/Mcp1 complexes are coordinated in space and time. One possibility is that they compete for the same physical space at the iMT plus tip. As Klp5/Klp6/Mcp1 concentration increases it may physically displace the Tea2/Tip1/Mal3 complex from the plus tip and induce MT catastrophe. An alternative, but not necessarily exclusive, possibility is that the Tea2/Tip1/Mal3 complex may occlude the Klp5/Klp6/Mcp1 complex from the plus tip until the iMT encounters the cell end. To test these models, we monitored the relative positions and levels of components of the Tea2/Tip1/Mal3 and Klp5/Klp6/Mcp1 complexes at the iMT plus tip. Importantly, dual imaging of *tea2-GFP tip1-tdTomato* cells shows that Tea2 and Tip1 occupy the same region of the MT plus tip and remain at the same intensity both before and after MT contact with the cell end cortex, consistent with the fact that these proteins are components of the same complex (Figure 3 - figure supplement 1A-B). By contrast, dual imaging of *klp5-mNG klp6-mNG tip1-tdTomato* cells reveals that Klp5/Klp6 accumulates behind Tip1 as the iMT grows and dwells at the cell end, as judged by maximal intensity measurements of the fluorescence (Figure 3A-B & Figure 3 - figure supplement 2). Importantly, the Tea2/Tip1/Mal3 and Klp5/Klp6/Mcp1 complexes converge at the plus tip approximately 10 seconds before MT shrinkage is detected. This behaviour is also observed in the absence of Mcp1, although in this case Klp5/Klp6 remains behind the Tea2/Tip1/Mal3 complex at the plus tip for longer as the MT dwells at the cell end (Figure 3C & Figure 3 - figure supplement 3).

**Figure 3.**
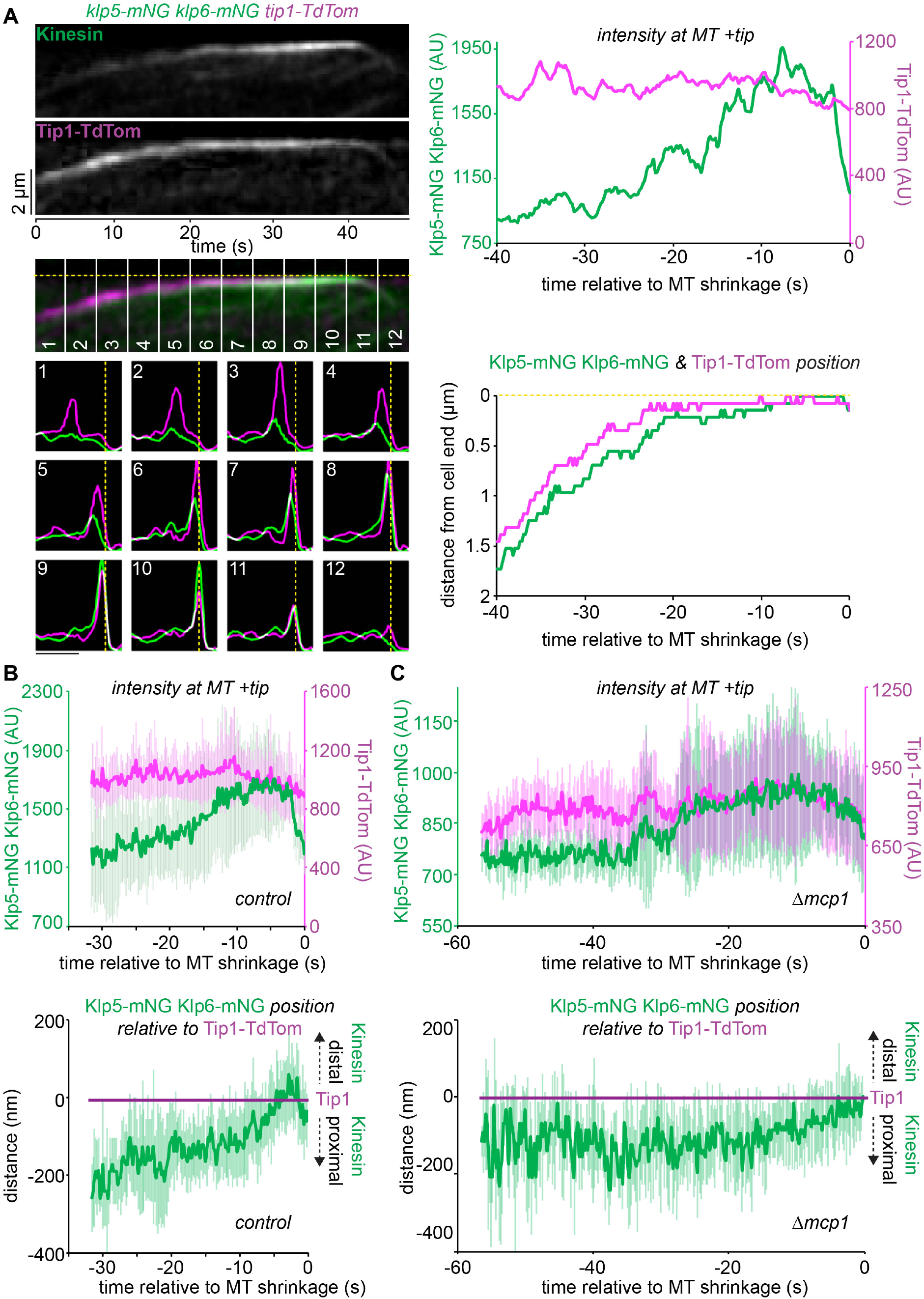
Klp5/Klp6 accumulates behind Tea2 at the microtubule plus tip until just prior to catastrophe. **(A)** A representative kymograph (top left panels) showing fluorescently-tagged Klp5/Klp6 (Kinesin) co-imaged with fluorescently-tagged Tip1 (Tip1-Tdtom). Dashed yellow line indicates the cell end. Plots (bottom left panels) representing both the fluorescence intensity and position of Klp5/Klp6 (green) and Tip1 (magenta) corresponding to the numbered sections of the kymograph. Bar, 2µm. Intensity of the maximal value pixel at the plus tip (top right panel) and its position relative to the cell end (bottom right panel) for Klp5/Klp6 (green) and Tip1 (magenta) relative to the time of microtubule shrinkage (seconds). **(B)** Data quantitated from multiple kymographs (n=7) of the type illustrated in (A) to display Klp5/Klp6 and Tip1 levels at the MT plus tip (top plot) and the position of Klp5/Klp6 relative to Tip1 (bottom plot) before MT shrinkage. Error bars show standard deviation. **(C)** Klp5/Klp6 is maintained behind Tip1 for longer in the absence of Mcp1. Data are quantitated from multiple kymographs (n=8) of the type illustrated in (Figure 3 - figure supplement 3) where fluorescently-tagged Klp5/Klp6 is co-imaged with fluorescently-tagged Tip1 in the absence of Mcp1 and presented as in (B).

These results indicate that the “antenna model” for MT age- and length-dependent accumulation of Kinesin-8 has validity but cannot solely explain the position-dependent interphase function of Klp5/Klp6 in fission yeast. The Klp6/Klp6/Mcp1 complex accumulates at MT plus tips over time, but Kinesin-8 walks only marginally faster than iMT polymerisation, so that its accumulation is largely dependent on MT growth slowing at cell ends rather than on MT length. More importantly, our data suggest a new model to explain iMT length control in fission yeast, in which the MT-stabilising Tea2/Tip1/Mal3 Kinesin complex binds ahead of the MT-destabilising Klp5/Klp6/Mcp1 complex and restricts its access to the iMT plus tip until the MT reaches the cell end (Figure 4). The earliest events in the sequence that results in MT catastrophe remain unclear, but we do see clearly that a slowing of iMT plus tip growth precedes any measurable change in the Tea2 complex signal. One possibility is that slowing down of iMT growth near cell ends allows Kinesin-8 to build up to a critical concentration that enables it to invade the space occupied by Tea2 and/or displace it from plus tips. Regardless, our work emphasises that iMT length control in cells is a complex emergent property reflecting competition and collaboration between multiple factors that influence tubulin dynamics. In the case of fission yeast, MT-destabilising Kinesin-8 motors accumulate at plus ends in a manner that is dependent on MT length and lifetime (as in the antenna model), but access to the MT plus end is controlled in time and space by an engagement mechanism that slows growth and favours unloading of the Tea2 complex. These two mechanisms act as separable layers of control that overlie the fundamental mechano-sensitive dynamics of tubulin itself at iMT plus ends.

**Figure 4.**
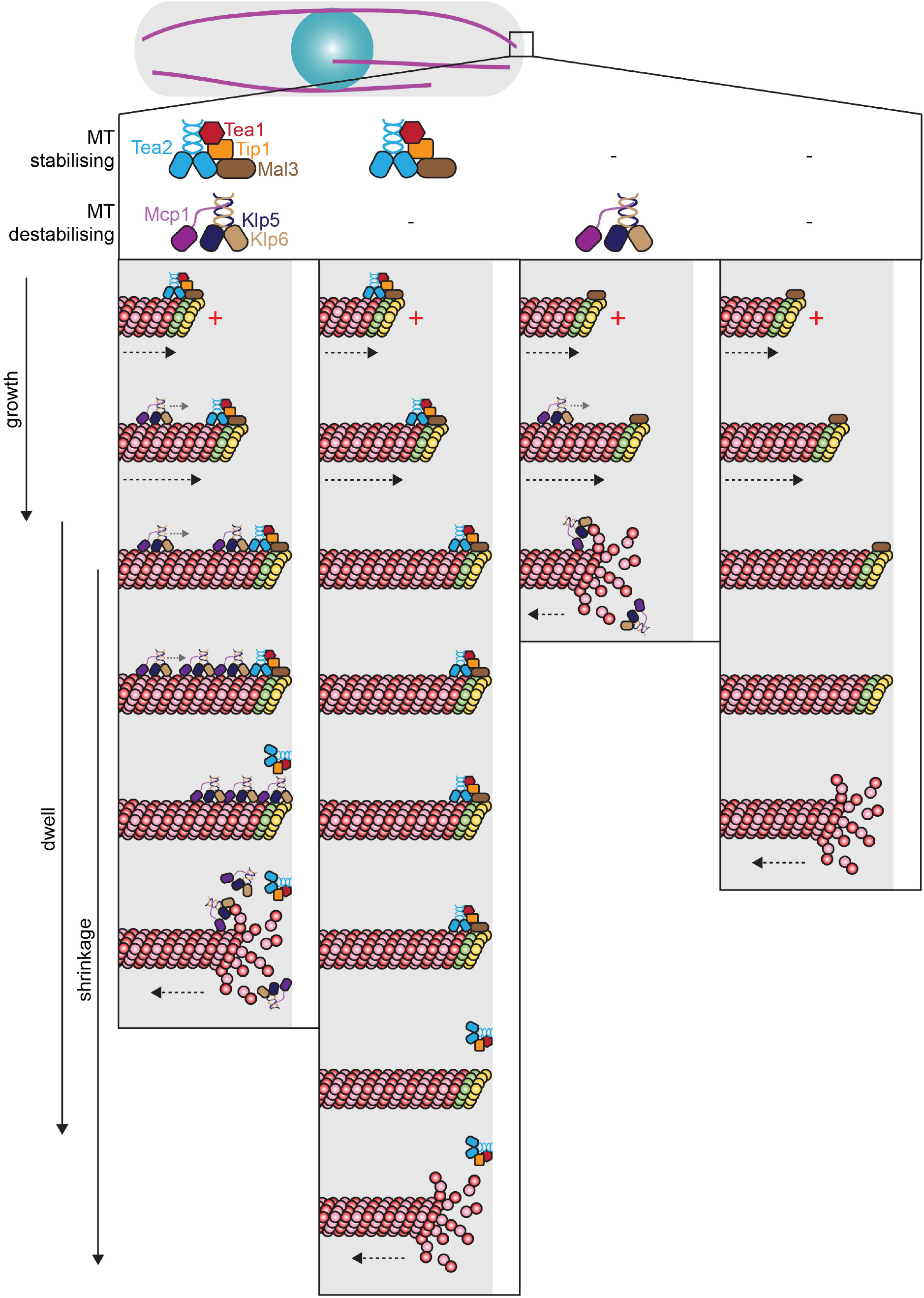
Destabilising Kinesin-8 accumulates behind the stabilising Tea2 kinesin complex to ensure microtubule disassembly at the cell end. The MT stabilising Tea2 complex (Tea2/Tip1/Mal3) binds to MT plus tips containing GTP-β-tubulin (green and yellow subunits) as they grow towards the cell end. The MT destabilising Kinesin-8 (Klp5/Klp6/Mcp1) complex walks along the MT lattice and accumulates behind the Tea2 complex, particularly as the MT dwells at the cell end, but is prevented from accessing the plus tip by the presence of Tea2. The Kinesin-8 complex may then either displace the Tea2 complex from the plus tip, triggering MT catastrophe, or initiate MT depolymerisation resulting in the loss of the Tea2 complex and the GTP cap. An alternative, but not necessarily exclusive, possibility is that displacement of Tea2 from the MT plus tip initiates Kinesin-8 mediated MT disassembly. In the absence of the MT destabilising Klp5/Klp6/Mcp1 complex (second column), MTs dwell for longer at the cell end and the Tea2 complex remains at the plus tip during this period. Klp5/Klp6/Mcp1 is therefore essential for regulating timely dissociation of the Tea2 complex from plus tips. In the absence of Tea2 (third column), Kinesin-8 accumulates at, and has unrestricted access to, the plus tip resulting in frequent MT catastrophe before reaching the cell end. When both stabilising and destabilising kinesin complexes are absent (fourth column) MTs reach and dwell at the cell end and may undergo catastrophe as a consequence of compressive force encountered at the cell end.

## Materials and Methods

### Cell Culture and Strain Construction

Media and the growth and maintenance of strains were as described previously (Moreno et al., 1991). All experiments were performed at 29.5°C unless stated. A full list of strains and oligonucleotides used in this study can be found in Supplementary File 1 and Supplementary File 2 respectively.

Deletion of *mcp1* with *natMX6*, *hphMX6* and *kanMX6*, C-terminal tagging of *mcp1* with *GFP*, and C-terminal tagging of *klp5* and *klp6* with *mNeonGreen* were carried out by two-step PCR gene targeting as described previously (Krawchuk & Wahls, 1999). The *mcp1*, *klp5* and *klp6* genes were also C-terminally tagged with RFP, mCherry or TdTomato. However, the fluorescence intensity of the resultant strains was too low and bleached too quickly to enable time-lapse live-cell imaging. N-terminally tagged *lys1::nmt1-GFP-mcp1* was constructed by amplifying the *mcp1* ORF flanked by *attB1* sites. The resulting PCR product was cloned into *pDONR221*^TM^ and then shuttled to *pLYS1U-HFG1c* using Gateway LR clonase II enzyme mix (Invitrogen, USA). Following linearisation with NruI this construct was integrated into *S. pombe* strains at the *lys1* locus and confirmed with PCR.

### Live-Cell Fluorescence Microscopy

Live-cell analyses were performed in imaging chambers (CoverWell PCI-2.5, Grace Bio-Labs) filled with 1 mL of 1% agarose in minimal medium and sealed with a 22 × 22 mm glass coverslip.

Figure 1B, Figure 1 - figure supplement 1A-C, Figure 2A, Figure 2E, and Figure 2 - figure supplement 2 used a single-camera Andor spinning disk confocal system comprising of a Nikon Ti-Eclipse microscope base equipped with a Plan Apo Vc 100x/1.40 N.A. objective lens, a Yokogawa CSU-XI confocal unit, an Andor iXon Ultra EMCCD camera with 2x magnification adapter and Andor iQ3 software. 488 nm and 561 nm solid state diode (power 50/100 mW) lasers were used for excitation. All images were processed and analysed using ImageJ (Fiji Life-Line version). Temperature was maintained at 29.5°C by a cage incubator controlled by OKO-touch (OKOLAB, Italy)

Figure 1C-F, Figure 1 - figure supplement 3F-G, Figure 2B-D,F, Figure 2 - figure supplement 1 & 3, Figure 3, Figure 3 - figure supplement 1-3 were generated using an Andor TuCam spinning disk confocal system comprising of a Nikon Ti-Eclipse microscope base equipped with a 100x/1.45 NA objective lens (Nikon CFI Plan Apo Lambda), a Nikon PFS (perfect focus system), a Yokogawa CSU-X confocal unit, and 2 Andor iXon Ultra EMCCD cameras with 2x magnification adapter and Andor TuCam two camera imaging adapter (beam splitter: Semrock DiR561). The final pixel size with 100x lens is 69 nm/pixel (measured). The light source is an Andor ILE laser unit equipped with solid state 50 mW lasers (488 & 561 nm). Temperature was maintained at 29.5°C by an OKO Lab hated enclosure.

### Registration of Images in the TuCam System

Registration of the 2 channels recorded simultaneously by 2 separate cameras was performed after acquisition using a reference image and ImageJ plugins for both image registration and transformation. Before every experiment, 3 2-channel reference images of 0.5 µm beads (calibration slide, Tetraspec Molecular Probes) in both the mNeonGreen/GFP and TdTomato/mCherry channels were captured. Following the experiment, reference images were again collected to compare for changes in calibration during the imaging session. Next, the channels in one of the reference images were registered using similarity transformation in the ImageJ plugin *RVSS* (Register Virtual Stack Slices) which performs registration using translation, rotation and isotropic scaling. This file was further tested on the remaining 2-channel reference images using the ImageJ plugin *TVSS* (Transform Virtual Stack Slices) and the transformation file saved for further use. Registered images were inspected to confirm correct alignment of both channels across the whole field of view. Finally, the transformation file was used to register the channels of the corresponding batch of experimental data.

### Kymograph Generation and Analysis

Kymographs were produced from data generated on the TuCam system. Mid-log phase cells were imaged for both green (mNeonGreen and GFP) and red (TdTomato and mCherry) tags simultaneously with 150 ms exposure every 200 ms for 2 minutes using 6% laser power for both channels. PFS was set to ON for all time points. Camera noise was suppressed by filtering all images using the Mean Filter in ImageJ set to 1×1. A single focal plane was used to increase temporal resolution. Only microtubules that remained in this plane throughout image acquisition were used to generate kymographs with the ImageJ plugin *KymoResliceWide* set to generate average maximum intensities for each frame perpendicular to a manually drawn line 10 pixels wide with spline fit.

Kymographs were analysed for fluorescence levels at the MT plus tip relative to MT length using a manually drawn line. MT shrinkage was defined as the first frame in which unambiguous MT shrinkage was first observed. MT growth speed was measured by calculating the gradient of mCherry through time during the portion of the kymograph where MTs were not in contact with the cortex. Klp5/Klp6 speed was measured by calculating the gradient of lines tracking mNeonGreen puncta on MT lattices. For the analyses presented in Figure 3 and Figure 3 - figure supplement 1 & 3 relating to maximal intensity pixel values, both channels from kymographs were saved as text images giving the intensity of every pixel for both channels. The pixel with maximal intensity was then determined for each channel at every time point and its position and fluorescence level recorded. The numbered plots that correspond to the numbered 20 pixels (4 s) wide segments of the kymographs show averaged fluorescence levels for each channel (relative to 100% maximum intensity for each trace) and position. These plots were generated using the *Gel Analyzer* tool in ImageJ.

### Relative Fluorescence Intensity Quantification

To compare the levels of the fluorescently-tagged proteins in two strain backgrounds, we performed mixing experiments to allow quantification within the same field of view on the TuCam system. Mid-log phase cells were excited for both green (mNeonGreen and GFP) and red (mCherry) tags simultaneously with 150 ms exposure for each of 11 Z sections (0.3 µm apart) every 2 s for 4 minutes using 6% laser power for both channels. PFS was initialised every 5 time points. Camera noise was suppressed by filtering all images using the Mean Filter in ImageJ set to 1×1 and Z stacks were flattened by maximum intensity projection. Fluorescence levels were quantified using a 5 x 5 pixel oval region of interest and determining the mean value. For plus tip values this signal had to be associated with the end of an iMT and was recorded in the frame immediately prior to MT shrinkage, for nuclear signals measurements were taken in the first frame. Background subtraction was performed against the cytoplasmic signal in all instances. Mitotic cells were excluded from the analysis.

### iMT Dwell Time Analysis

Cells expressing *GFP-atb2* were grown until mid-log phase and then microtubules imaged live using the single-camera Andor spinning disk confocal system. 8 z-stacks (0.5 μm apart) were taken with exposure times of 100 ms every 5 s for 10 minutes for GFP. These time-lapse series were then flattened in the Z dimension by maximum intensity projection in ImageJ. Dwell time was defined as the length of time the iMT plus end remained in the final 1.1 μm of a cell, with the cell perimeter defined by background GFP fluorescence. 1.1 μm is the mean length (plus one standard deviation) for the curved end of *wild type* cells and accounts for 84% of shrinkage events. Dwell times were measured for the full 10 minutes of each movie. ∼100 dwell times were measured for each condition.

### Imaging of plus tip proteins relative to iMT length

For Figure 1 - figure supplement 1A-C and Figure 2A, mid-log phase cells expressing fluorescently-tagged Klp5, Klp6, Tea2 or Mcp1, together with fluorescently-tagged MTs, were imaged live on the single-camera Andor system. 8 z-stacks (0.5 μm apart) were taken with green fluorophores (GFP and mNeonGreen) and red fluorophores (mCherry) excited for 200 ms every 4.8 s for no more than 5 minutes. Movement of Mcp1 puncta over time (Figure 1 – figure supplement 1B-C) were tracked using the ImageJ plugin *MTrackJ*. Klp5/Klp6 and Tea2 levels at plus tips relative to MT length (Figure 2A) were analysed manually with mNeonGreen and GFP levels background substituted and normalised against the mCherry mean intensity of iMTs. iMT length was measured by using the line tool in ImageJ with the anti-parallel MT bundle used to define the minus end. Measurements were performed over 3 independent experiments.

### Measurement of the Pre-anaphase Mitotic Index

Mid-log phase *fta3-GFP sid4-TdTomato*, or *cdc13-GFP sid4-TdTomato* cells were grown at 30°C and, following fixation in 3.7% formaldehyde for 10 minutes, mounted in medium containing DAPI to label DNA. Stacks of 18 z sections (0.2 μm apart) were taken on a wide field microscope system using a Nikon TE-2000 base with a 100x/1.49 N.A. objective lens equipped with a Photometrics Coolsnap-HQ2 liquid cooled CCD) camera and analysed using MetaMorph (version 7.5.2.0, MAG, Biosystems Software). The percentage of cells with Cdc13-GFP on the spindle pole bodies and mitotic spindle or the percentage of cells with kinetochores between separated poles prior to anaphase was determined. Exposure times of 1 s were used for GFP and TdTomato and 0.25 s for DAPI. For each experiment, at least 500 cells were counted, and each experiment was conducted three times.

### Analysis of Cell Curvature

Since polarity defects are emphasised in longer cells, *cdc25-22* cells were grown at 28°C until mid-log phase at which point they were shifted to 35.5°C for 6 hours to arrest at G2/M and then imaged live on the Nikon wide field system. 18 z-stacks (0.2 µm apart) were taken with a 100 ms exposure time for bright-field. Each experiment analysed at least 280 cells, was repeated 3 times and excluded cells that were less than 14 µm long. The z-stack containing the middle of the cell was used for analysis, whereby a line was drawn manually between the two cell ends in the middle of the cell using the line tool in ImageJ. The length of this line (L, length) was then compared to that of a straight line between the cell ends (E, Euclidean distance) using the measure function of ImageJ. The ratio between these values, converted to a percentage, gives a measure of the curvature of each cell. The distribution of cell lengths was unaltered between backgrounds.

### Biochemistry

Cells were lysed in buffer containing 50 mM HEPES (pH 7.6), 75 mM KCl, 1 mM MgCl_2_, 1 mM EGTA, 0.1% Triton X-100, 1 mM DTT, 1 mM PMSF and complete EDTA-free protease inhibitor cocktail (Roche). Protein concentration was quantified by Bradford assay (Biorad, USA). For co-immunoprecipitation, 2 mg protein extract was incubated with 5 μL (∼ 5 μg) rabbit *α*GFP (Immune Systems) (Immune sera, I) or 2.5 µL (∼5 μg) normal rabbit serum (Pre-Immune sera, PI) for 30 minutes and subsequently with 20 µL protein A-sepharose beads (GE Healthcare, USA) for 45 minutes. Following centrifugation at 4°C, beads were washed 3 times in lysis buffer before being heated at 100°C in SDS-sample buffer for 6 minutes.

Precipitates using immune (I) or pre-immune (PI) sera and whole cell extracts (WCE) were migrated using SDS-PAGE and transferred to nitrocellulose membrane. GFP was detected using anti-GFP sheep polyclonal (1/5000, Kevin Hardwick) and HRP-conjugated anti-sheep secondary antibody (1/10000, Jackson ImmunoResearch, USA). Myc was detected using anti-Myc mouse monoclonal (1/500, Abcam, UK) and HRP-conjugated anti-mouse secondary antibody (1/10000, GE Health Care, USA). Tubulin was detected by anti-TAT1 mouse monoclonal (1/1000, Keith Gull) and HRP-conjugated anti-mouse secondary antibody (1/10000, GE Health Care, USA). ECL detection was performed using Amersham ECL system (GE Healthcare).

### Statistics

Since some data sets were not normally distributed, as ascertained by the Shapiro-Wilk test, the nonparametric two-sample Kolmogorov-Smirnov test was used throughout to determine the probability that two data-sets come from different distributions. Text files were manipulated in Excel (version 15.33, Microsoft) and data were analysed by R (version 3.3.3, R Core Team) and RStudio (version 1.0.136, RStudio, Inc).

## Acknowledgements

The authors are grateful to Kevin Hardwick and Keith Gull for their gifts of antibodies and to Fred Chang, Ken Sawin, Takashi Toda, Xiangwei He, Mitsuhiro Yanagida, Jean-Paul Javerzat and Paul Nurse for providing fission yeast strains.

---

**Figure 1 - figure supplement 1.**
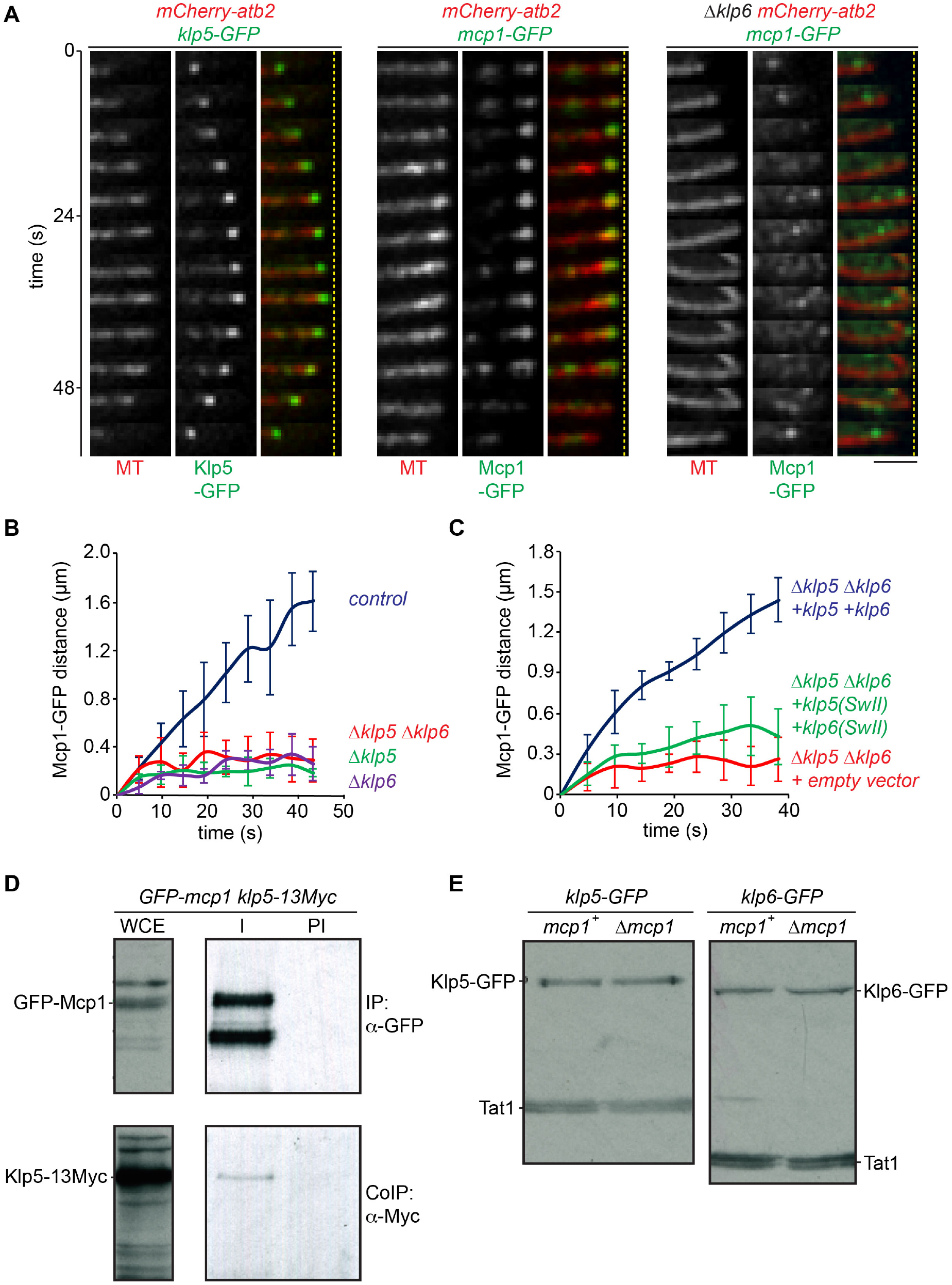
Mcp1 interacts with Klp5/Klp6 and requires its motor activity to accumulate at microtubule plus tips. **(A)** Accumulation of Mcp1 at the microtubule plus tip depends on Klp6. Panels showing fluorescently-tagged microtubules (MT) and either Klp5-GFP (left panel), Mcp1-GFP (middle panel) or Mcp1-GFP in the absence of Klp6 (right panel) imaged every 4.8 seconds. Dashed yellow lines indicate the cell end. Bar, 5µm. **(B)** Mcp1 movement depends on Klp5/Klp6. Plot shows the mean distance moved over time of five GFP iMT-associated puncta from each of the indicated backgrounds. Error bars show standard deviation. **(C)** Mcp1 movement depends on Klp5/Klp6 motor activity. Plot shows the mean distance moved over time of five GFP iMT-associated puncta from each of the indicated backgrounds. Error bars show standard deviation. **(D)** Mcp1 interacts weakly with Klp5. Log phase cultures of *GFP-mcp1 klp5-13myc* cells were harvested and lysed. Proteins were immunoprecipitated from 2mg of whole cell extract (WCE) using rabbit α-GFP antibodies (I) or pre-immune control (PI), migrated on an SDS-PAGE gel and probed with either sheep α-GFP or mouse α-Myc antibodies. 50 μg of WCE was run for comparison. **(E)** Cellular levels of Klp5/Klp6 are not altered by the absence of Mcp1. Log phase cultures of cells expressing *klp5-GFP* (left panels) or *klp6-GFP* (right panel) in control or *Δmcp1* cells were lysed and proteins extracted. 50 µg of each was then migrated by SDS-PAGE, transferred to nitrocellulose membrane and probed with both *α*-GFP to determine protein level and α-Tat1 to use tubulin as a loading control.

**Figure 1 - figure supplement 2.**
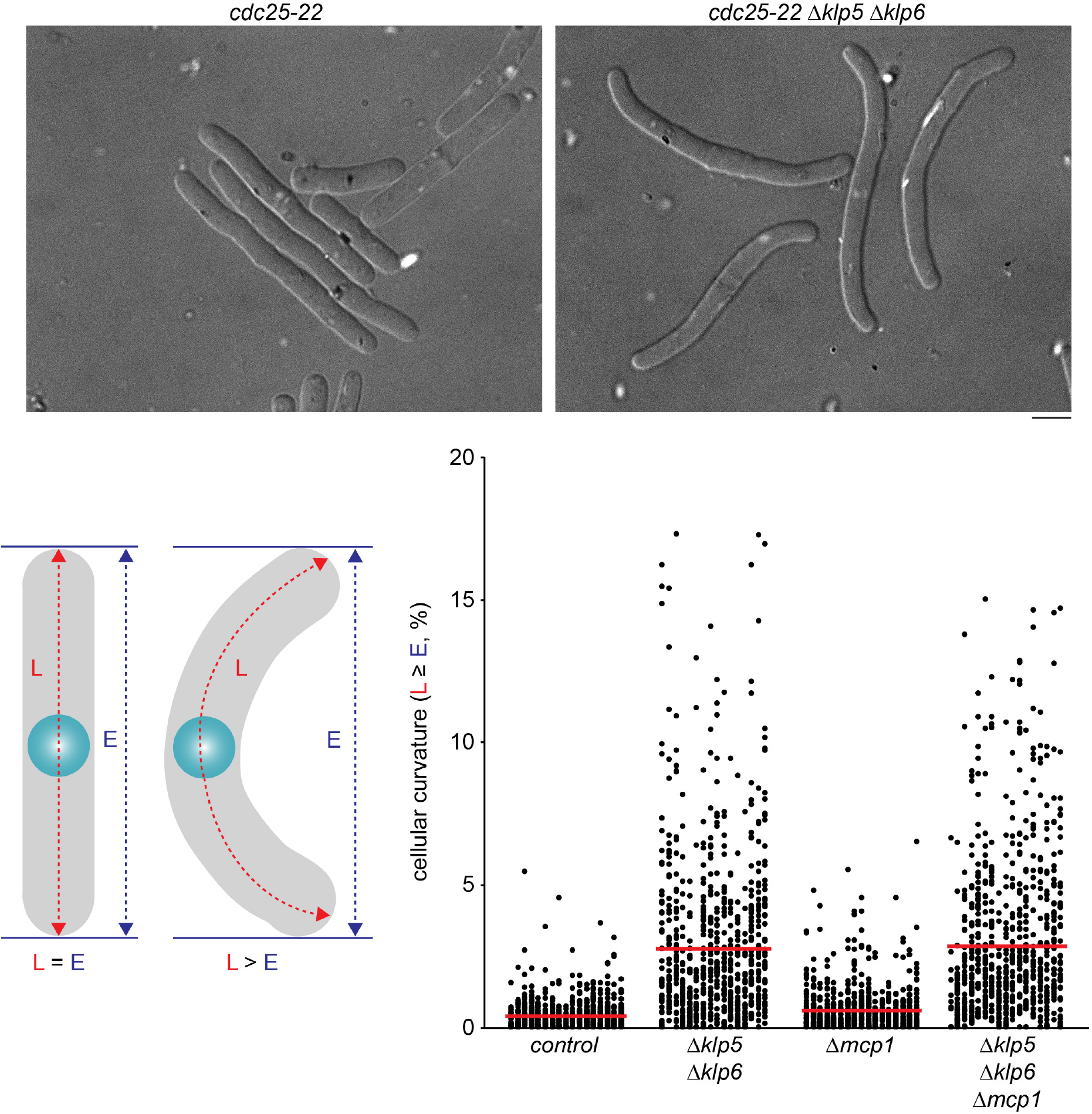
Mcp1 is dispensable for cellular polarity control Images show representative *cdc25-22* cells (left panel) or *cdc25-22 Δklp5 Δklp6* cells (right panel) arrested at the restrictive temperature (35.5°C) for six hours. Bar, 5µm. Cellular curvature was quantitated, as in the schematic, by measuring both the cell length (length, L) and the distance between cell ends (Euclidean distance, E) and then calculating the ratio (L:E). These ratios, converted to percentages, are displayed on the plot, with red lines showing the mean value (∼850 cells were measured for each strain).

**Figure 1 - figure supplement 3.**
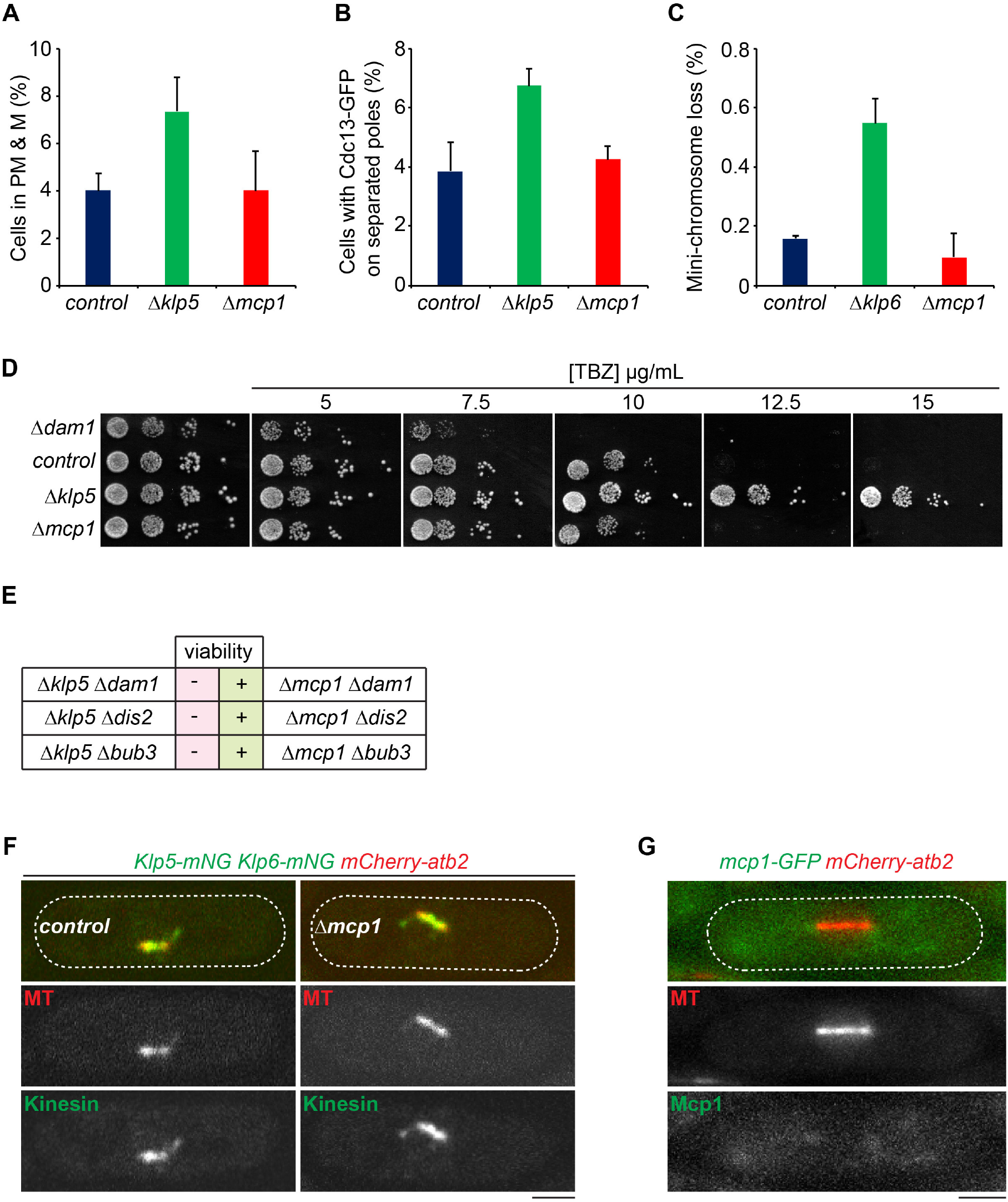
Mcp1 does not control the mitotic functions of Klp5/Klp6. **(A)** The pre-anaphase mitotic index is unaltered in the absence of Mcp1. Log phase cultures of control, *Δklp5* or *Δmcp1* cells expressing fluorescently-tagged kinetochore (Fta3) and spindle pole body (Sid4) proteins were imaged. The proportion of pre-anaphase mitotic cells with unseparated kinetochore pairs between poles was determined. The average of 3 independent experiments is plotted, error bars show standard deviation. **(B)** The timing of Cyclin B destruction is unaltered in cells lacking Mcp1. Log phase cultures of control, *Δklp5* or *Δmcp1* cells expressing fluorescently-tagged Cyclin B (Cdc13) and Sid4 were imaged. The proportion of cells with Cdc13-GFP on separated poles and spindles was determined. The average of 3 independent experiments is plotted, error bars show standard deviation. **(C)** Cells lacking Mcp1 are proficient at maintaining a mini-chromosome. Log phase cultures of control, *Δklp5* or *Δmcp1* cells expressing *ade6-M210* and carrying the *Ch16(ade6-M216)* mini-chromosome were grown in media lacking adenine and then individual cells plated onto media with minimal adenine. After 3 days, the proportion of cells that had formed colonies that were ≥ 50% red sectored, having lost the mini-chromosome in the first mitotic division, were scored as a proportion of all colonies. The average of 3 independent experiments is plotted, error bars show standard deviation. **(D)** The absence of Mcp1 does not alter sensitivity to thiabendazole. Log phase cultures of control, *Δklp5, Δdam1* or *Δmcp1* cells were serially diluted onto plates containing the indicated concentrations of the MT poison thiabendazole (TBZ) and grown for 3 days. Cells lacking Klp5 have elevated resistance to TBZ whereas cells lacking Dam1, a component of the DASH complex, display enhanced sensitivity to TBZ. **(E)** Strains lacking Dam1, Dis2 and Bub3 can survive without Mcp1 but not in the absence of Klp5 **(F)** Mcp1 does not influence localisation of Klp5/Klp6 to kinetochores and the spindle during mitosis. Representative images of mitotic cells expressing fluorescently-tagged MTs and Klp5/Klp6 in the presence (left panels) or absence (right panels) of Mcp1. Bar, 2µm. **(G)** Mcp1 does not localise to kinetochores or the spindle in mitotic cells. A mitotic cell expressing fluorescently-tagged MTs and Mcp1. Bar, 2µm.

**Figure 2 - figure supplement 1.**
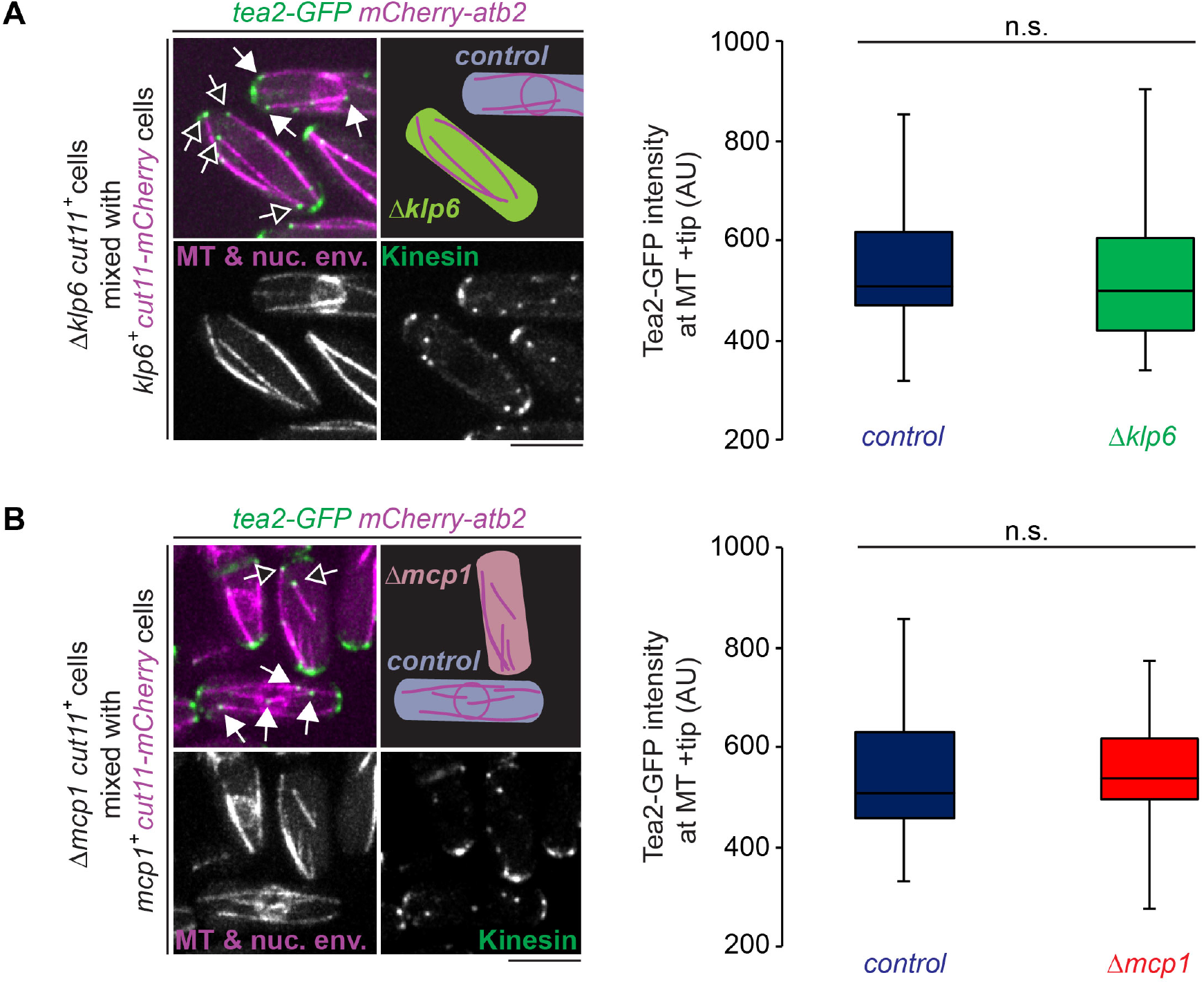
Klp5/Klp6/Mcp1 does not influence accumulation of Tea2 at microtubule plus tips. **(A)** The left panel shows a mixing experiment used to compare the intensity of fluorescently-tagged Tea2 on microtubule plus tips in either control (blue, closed arrowheads) or *Δklp6* cells (green, open arrowheads) in the same field of view. Control cells can be distinguished from *Δklp6* cells as they express a fluorescently-tagged nuclear envelope protein Cut11. Bar, 5µm. The box plot shows data quantitated from these experiments. Fluorescence values for Tea2-GFP in control (n=49) and *Δklp6* (n=50) cells were recorded in the frame prior to MT shrinkage. Measurements were normalised against cytoplasmic Tea2 levels in all cases. n.s. (non-significant) represents a p-value >0.05 between data sets calculated using the Kolmogorov-Smirnov test. **(B)** Shows the same experiment and layout as in (A) to compare Tea2 levels on microtubule plus tips in either control (blue, closed arrowheads) or *Δmcp1* (red, open arrowheads) cells in the same field of view. Fluorescence values for Tea2-GFP in control (n=50) and *Δmcp1* (n=50) cells were recorded in the frame prior to MT shrinkage.

**Figure 2 - figure supplement 2.**
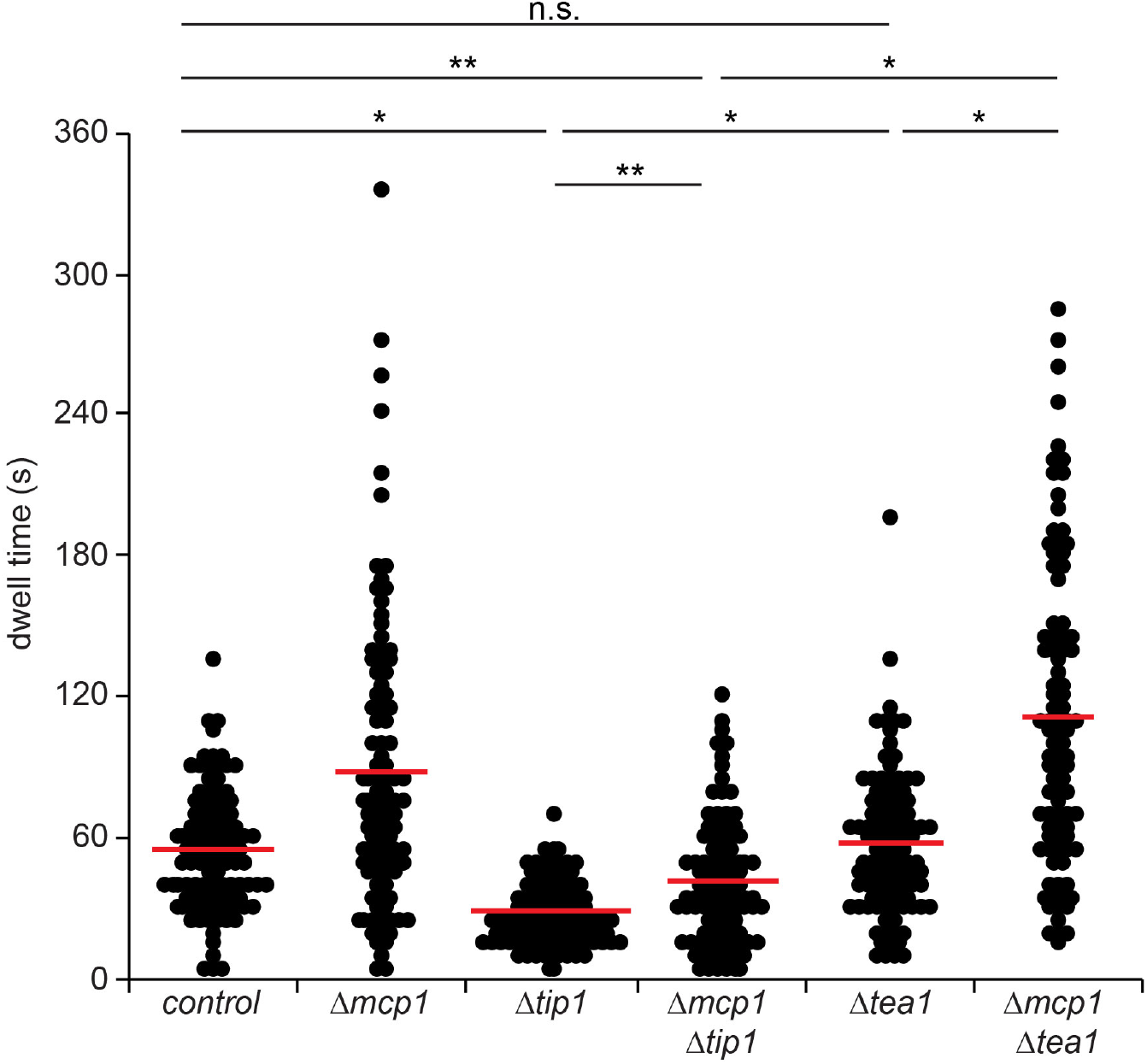
Tip1 stabilises microtubules in opposition to Mcp1 Cells expressing fluorescently-tagged tubulin were imaged every 5 seconds and the dwell time of 100 individual iMTs for each condition recorded within the final 1.1 µm of the cell. Red bars signify the mean. Single asterisks represent p-values <0.001, double asterisks <0.01 and n.s. >0.05 (non-significant) between data sets calculated using the Kolmogorov-Smirnov test.

**Figure 2 - figure supplement 3.**
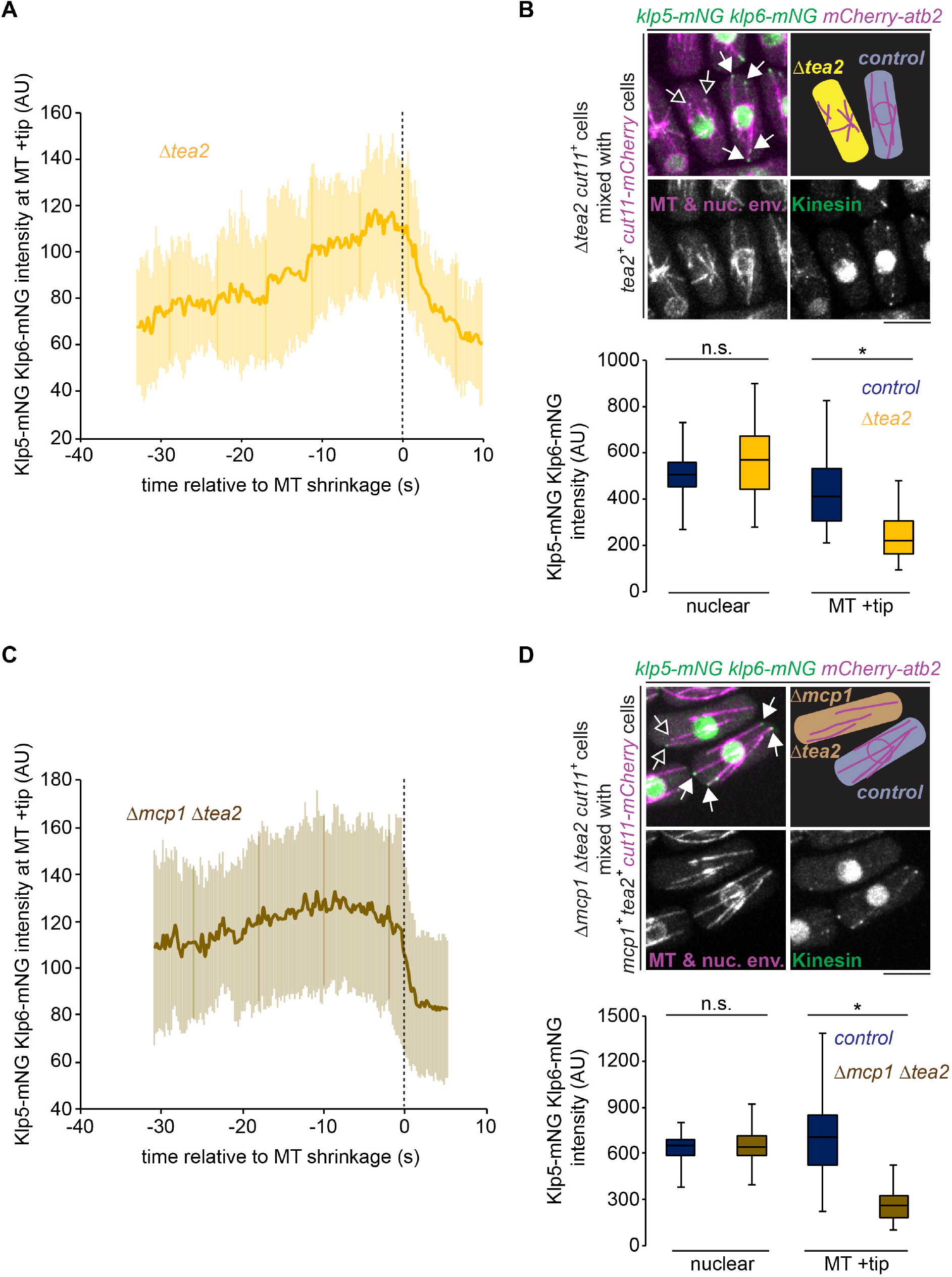
Absence of Tea2 or both Tea2 and Mcp1 alters plus tip Klp5/Klp6 levels at microtubule shrinkage. **(A)** Intensity of Klp5/Klp6 at microtubule plus tips relative to the time of microtubule shrinkage was quantified from multiple kymographs (n=16) of *Δtea2* cells. Error bars show standard deviation. **(B)** The top panel shows the mixing experiment used to compare fluorescently-tagged Klp5/Klp6 levels between control (blue, closed arrowheads) or *Δtea2* (yellow, open arrowheads) cells in the same field of view. Control cells can be distinguished from *Δtea2* cells as they also express a fluorescently-tagged nuclear envelope protein Cut11. Bar, 5µm. The box plot below shows data quantitated from these experiments. Fluorescence values were recorded in the first frame for nuclear levels of Klp5/Klp6 in control (n=37) and *Δtea2* (n=37) cells or levels of Klp5/Klp6 at the plus tip in the frame prior to microtubule shrinkage in control (n=37) and *Δtea2* (n=37) cells. Measurements were normalised against cytoplasmic Klp5/Klp6 levels in all cases. Asterisk represents a p-value <0.001 and n.s. >0.05 (non-significant) between data sets calculated using the Kolmogorov-Smirnov test. **(C)** Intensity of Klp5/Klp6 at microtubule plus tips relative to the time of microtubule shrinkage was quantified from multiple kymographs of *Δmcp1 Δtea2* cells (n=24). Error bars show standard deviation. **(D)** Shows the same experiment and layout as in (B) but with Klp5/Klp6 levels compared between control (blue, closed arrowheads) or *Δmcp1 Δtea2* (brown, open arrowheads) cells in the same field of view. Fluorescence values were recorded in the first frame for nuclear levels of Klp5/Klp6 in control (n=50) and *Δmcp1 Δtea2* (n=50) cells or levels of Klp5/Klp6 at the plus tip in the frame prior to microtubule shrinkage in control (n=50) and *Δmcp1 Δtea2* (n=50) cells.

**Figure 3 - figure supplement 1.**
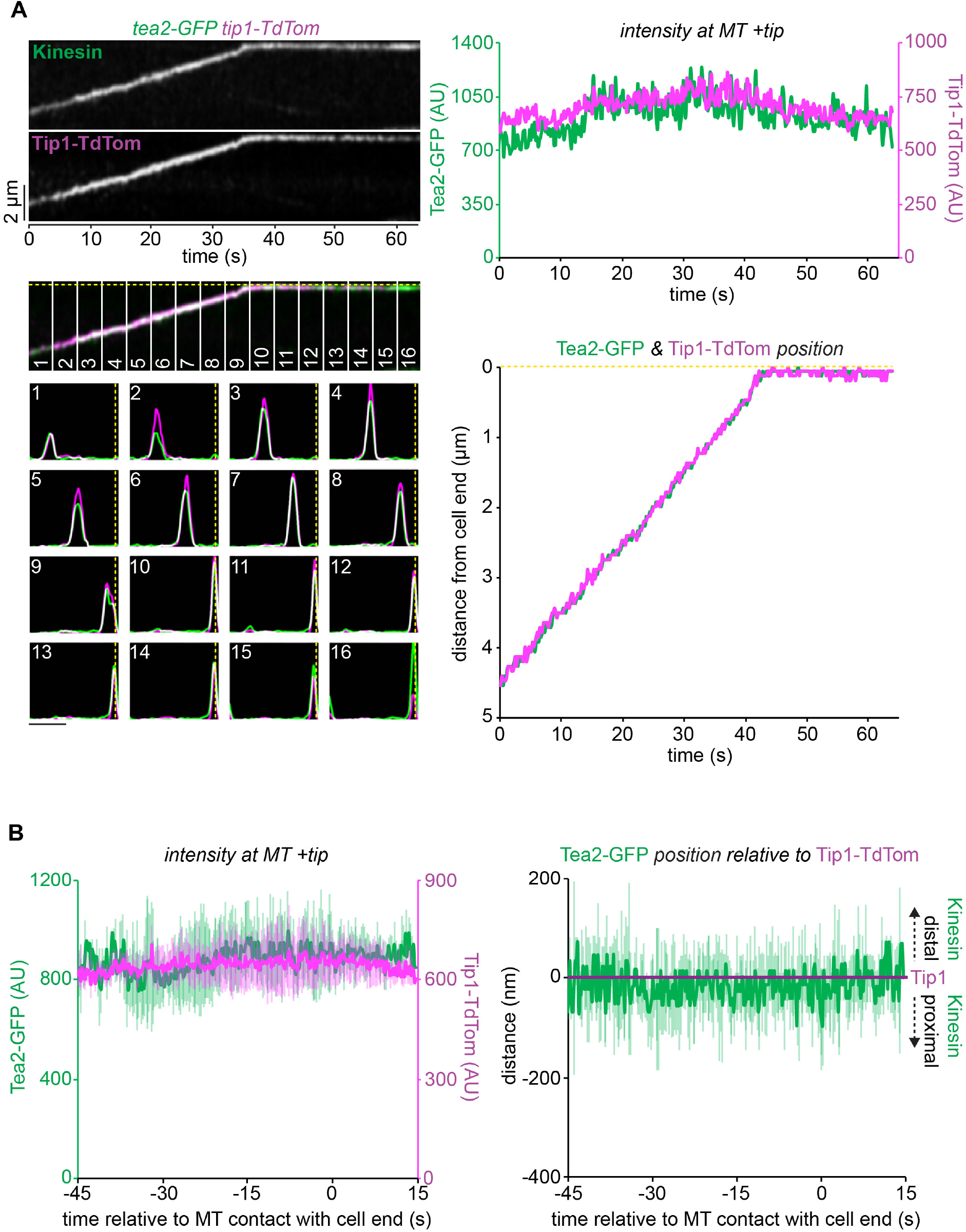
Tea2 and Tip1 co-localise at the microtubule plus tip A representative kymograph (top left panels) showing fluorescently-tagged Tea2 (Kinesin) co-imaged with fluorescently-tagged Tip1 (Tip1-Tdtom). Dashed yellow line indicates the position of the cell end. Plots (bottom left panels) representing both the relative fluorescence intensity and position for Tea2 (green) and Tip1 (magenta) corresponding to the numbered sections of the kymograph. Bar 2µm. Data quantitated from the kymograph by extracting the maximal intensity pixel value at the MT plus tip for Tea2 (green) and Tip1 (magenta) over time (top right) and the distance of these pixels from the cell end (bottom right). **(B)** Data quantitated from multiple kymographs (n=5) of the type illustrated in (A) to display Tea2 and Tip intensity at the MT plus tip (left plot) and the position of Tea2 relative to Tip1 (right panel) and relative to initial MT contact with the cell end. Error bars show standard deviation.

**Figure 3 - figure supplement 2.**
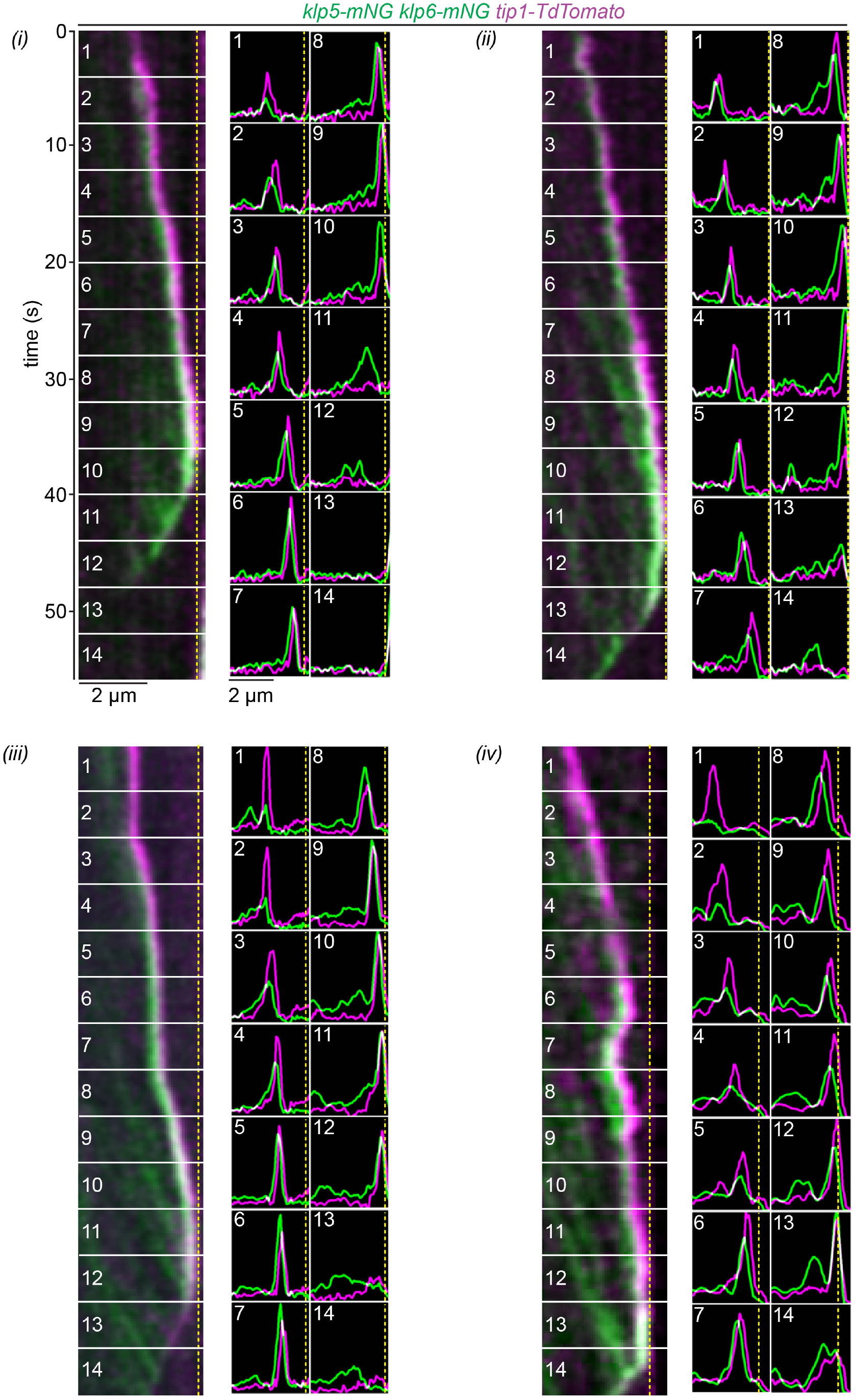
The relative intensity and location of Klp5/Klp6 and the T-complex characterises microtubule growth, dwell and shrinkage Kymographs of four additional movies (*i-iv*) of *klp5-mNeongreen klp6-mNeongreen tip1-Tdtomato* cells expressing fluorescently-tagged Klp5/Klp6 (green) co-imaged with fluorescently-tagged Tip1 (magenta). Dashed yellow line indicates the cell end. The associated plots show both the relative fluorescence intensity and position for Klp5/Klp6 and Tip1 corresponding to the numbered sections of the kymographs.

**Figure 3 - figure supplement 3.**
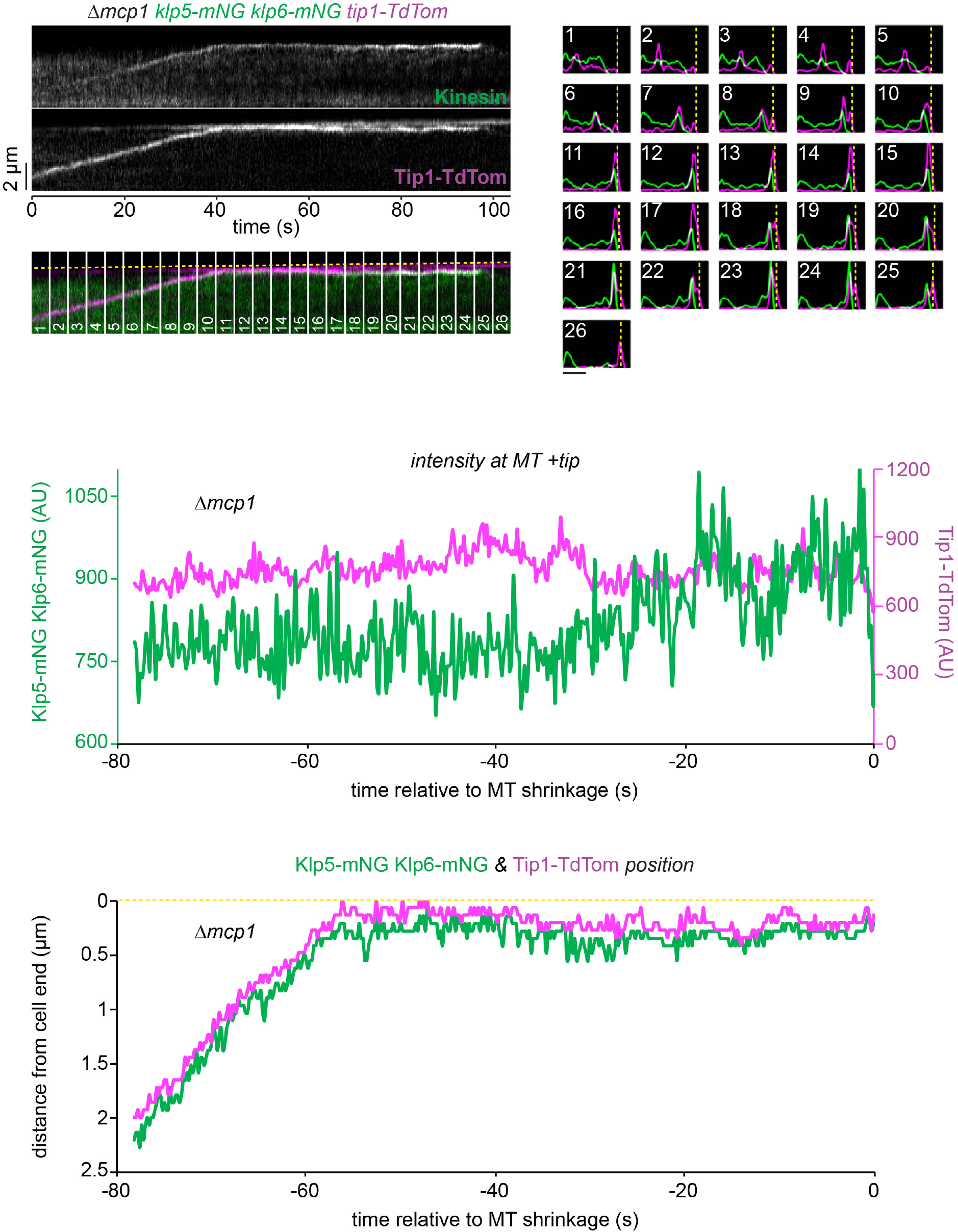
Klp5/Klp6 queues behind Tea2 for longer in the absence of Mcp1. A representative kymograph (top left panels) showing fluorescently-tagged Klp5/Klp6 (Kinesin) co-imaged with fluorescently-tagged Tip1 (Tip1-Tdtom) in the absence of Mcp1. Dashed yellow line indicates the position of the cell end. Plots (top right panels) showing both the relative fluorescence intensity and position for Klp5/Klp6 and Tip1 corresponding to the numbered sections of the kymograph. Bar, 2µm. Data were quantitated from the kymograph by extracting the maximal intensity pixel value at the MT plus tip over time for Klp5/Klp6 and Tip1 (middle panel) and by plotting the distance that these pixels are from the cell end relative to the time of microtubule shrinkage (bottom panel).

## Supplementary File 1. List of strains used in this study

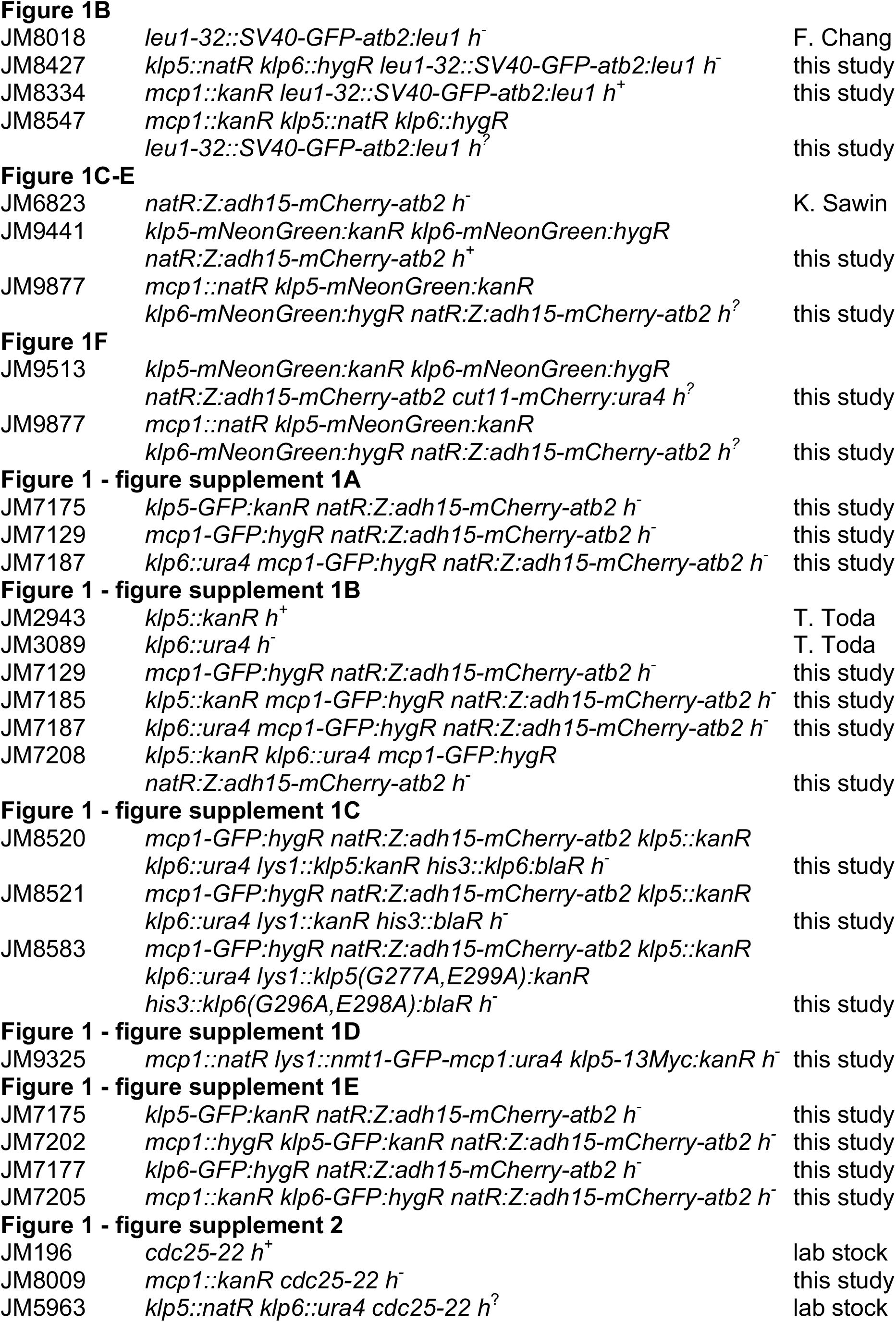
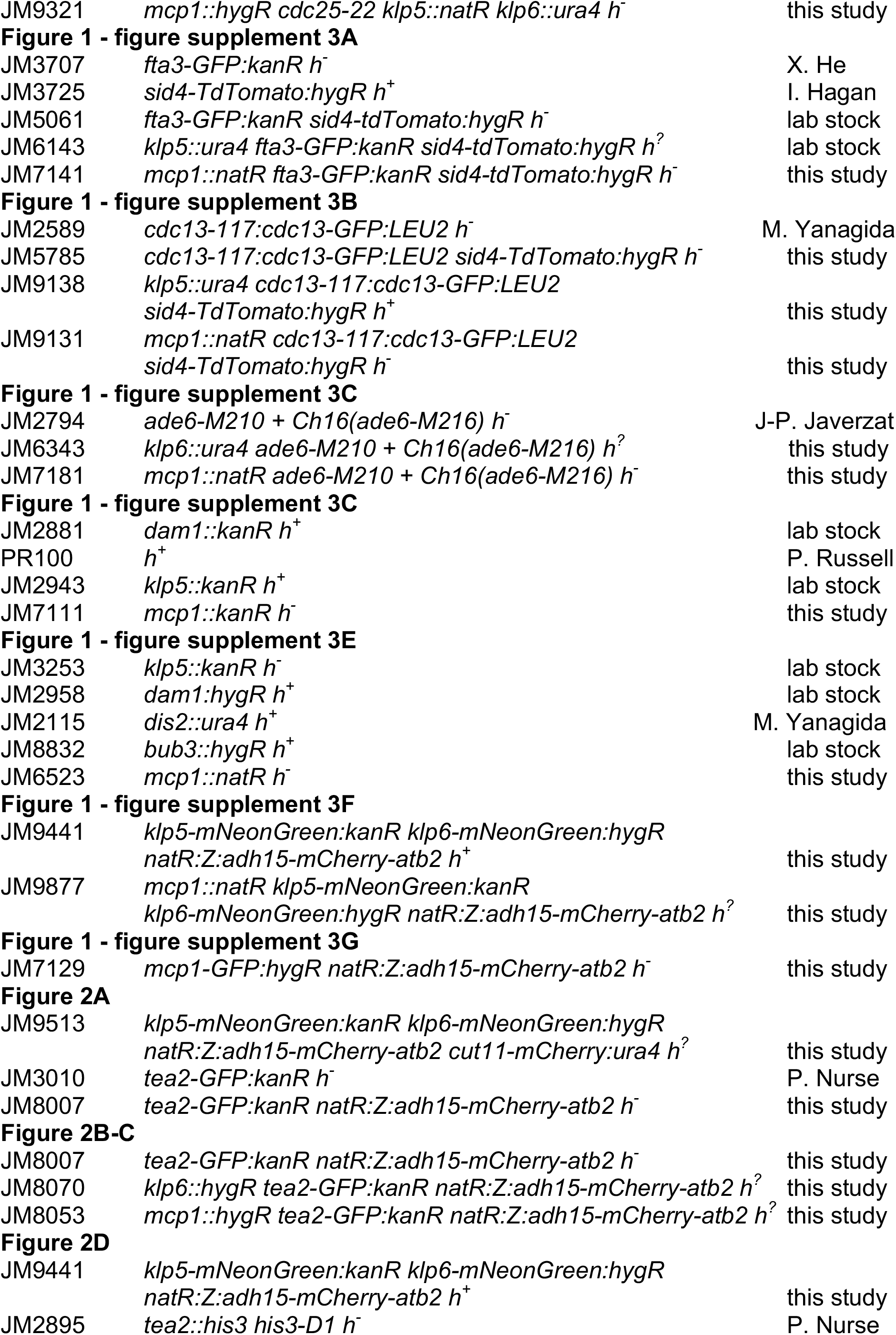
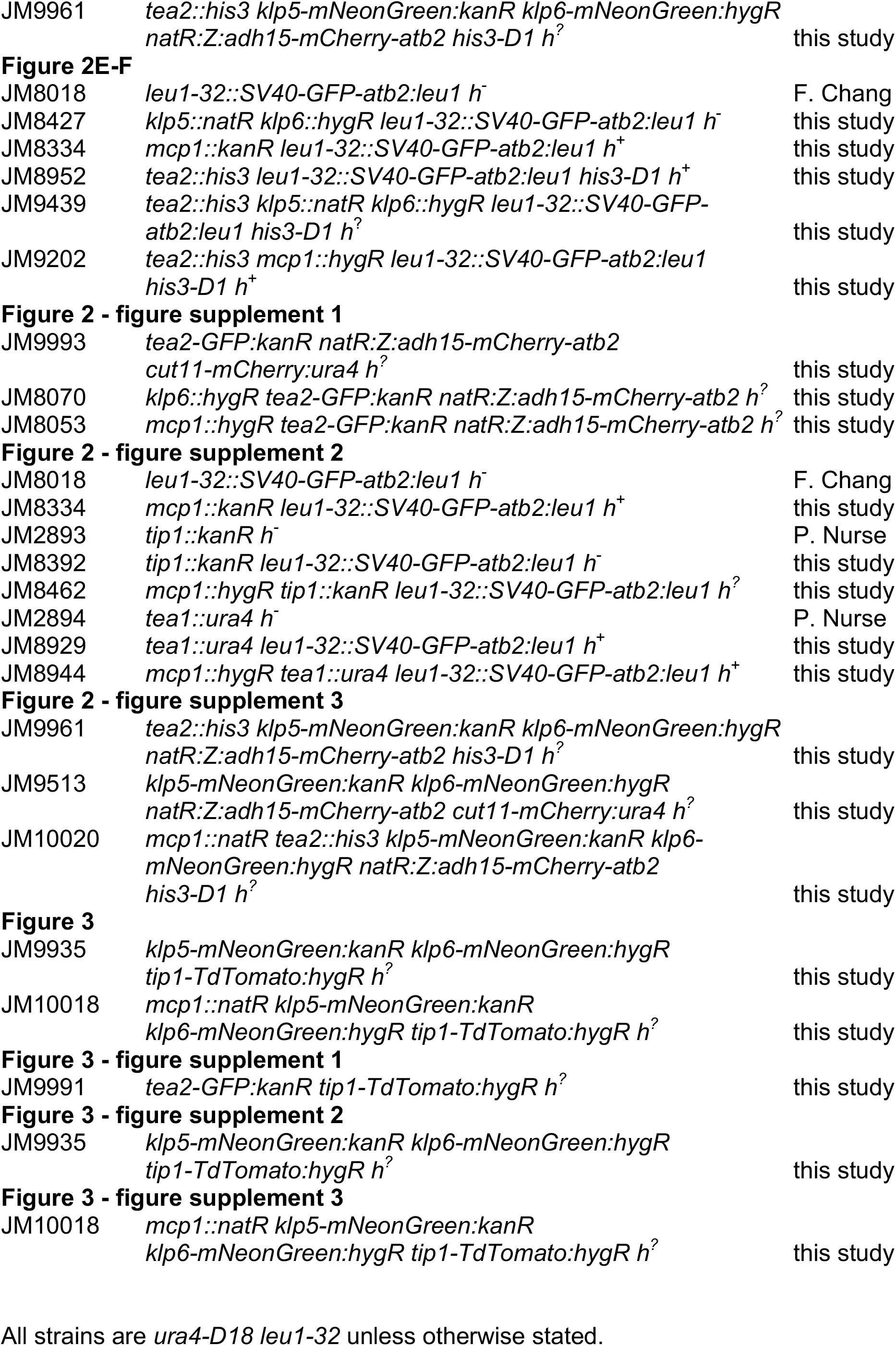

## Supplementary File 2. List of oligonucleotides used in this study

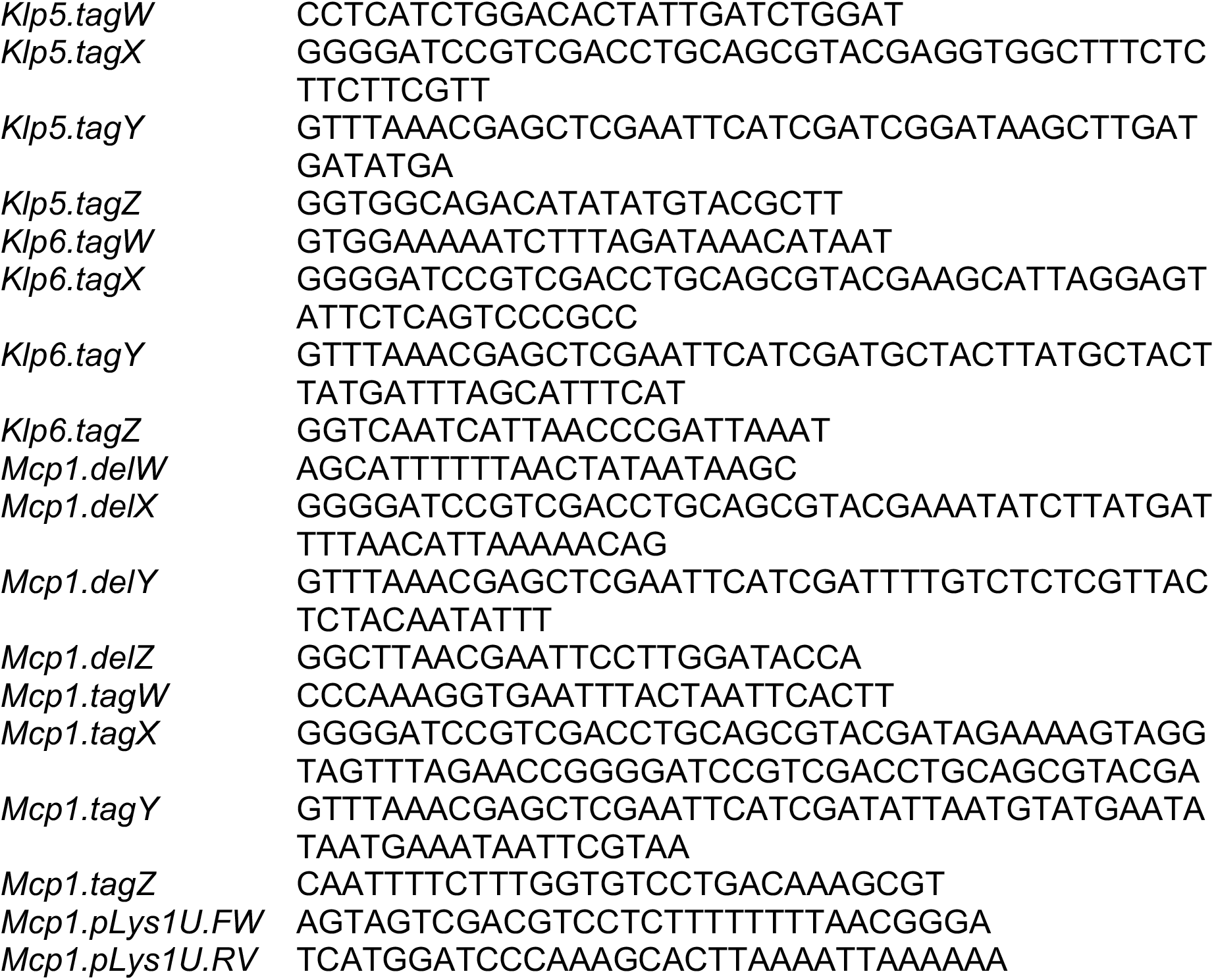

